# Dissection-independent production of a protective whole-sporozoite malaria vaccine

**DOI:** 10.1101/2020.06.22.164756

**Authors:** Joshua Blight, Katarzyna A. Sala, Erwan Atcheson, Holger Kramer, Aadil El-Turabi, Eliana Real, Farah A. Dahalan, Paulo Bettencourt, Emma Dickinson, Eduardo Alves, Ahmed M. Salman, Chris J. Janse, Frances Ashcroft, Adrian V. S. Hill, Arturo Reyes-Sandoval, Andrew M. Blagborough, Jake Baum

## Abstract

Complete protection against human malaria challenge has been achieved using infected mosquitoes as the delivery route for immunization with *Plasmodium* parasites. Strategies seeking to replicate this efficacy with either a manufactured whole-parasite or subunit vaccine, however, have shown only limited success. A major roadblock to whole parasite vaccine progress and understanding of the human infective sporozoite form in general, is reliance on manual dissection for parasite isolation from infected mosquitoes. We report here the development of a four-step process based on whole mosquito homogenization, slurry and density-gradient filtration, combined with free-flow electrophoresis that is able to rapidly produce a pure, aseptic sporozoite inoculum from hundreds of mosquitoes. Murine *P. berghei* or human-infective *P. falciparum* sporozoites produced in this way are 2-3-fold more infective with *in vitro* hepatocytes and can confer sterile protection when immunized intravenously with subsequent challenge using a mouse malaria model. Critically, we can also demonstrate for the first time 60-70% protection when the same parasites are administered via intramuscular (i.m.) route. In developing a process amenable to industrialisation and demonstrating efficacy by i.m. route these data represent a major advancement in capacity to produce a whole parasite malaria vaccine at scale.

**One-Sentence Summary:** A four-step process for isolating pure infective malaria parasite sporozoites at scale from homogenized whole mosquitoes, independent of manual dissection, is able to produce a whole parasite vaccine inoculum that confers sterilizing protection.

## Main Text

A vaccine against malaria is still desperately required to tackle the nearly half a million deaths caused by the disease each year [1]. To date, two distinct approaches to develop an effective malaria vaccine have been widely utilised, based either on the recombinant production and immunisation of dominant surface antigens of the *Plasmodium* parasite, or the labour-intensive production, purification and administration of live attenuated parasites. Despite decades of investment, however, current vaccines using either strategy have shown limited efficacy in reducing malaria infection in field trials [2,3].

The most advanced malaria vaccines within the development pipeline have focussed on the liver stages of parasite infection, specifically the infectious sporozoite form, injected by the mosquito during a blood feed [2,3]. Most have been based on either the dominant sporozoite surface antigen, circumsporozoite protein (CSP) [2] or on live-attenuated sporozoites themselves. CSP has been the subject of intensive investigation for more than forty years. Its most recent formulation within the RTS,S vaccine, confers moderate protection following challenge in phase III trials, with uncertain longterm efficacy [3]. Live-attenuated sporozoite vaccines [4–8] represent an entirely different strategy, arresting prematurely in hepatocytes [9,10], allowing both humoral and cellular immunity to develop offering long-term protection when delivered as a vaccine [6,11,12]. The most advanced of these, PfSPZ [13], has conferred sterile protection (complete protection) under controlled clinical conditions, but only when delivered intravenously, at high doses (270,000 sporozoites) and with multiple rounds of immunization (typically one prime and three boosts) [6,9–12,14]. PfSPZ has, like RTS,S, shown moderate efficacy in field trials [15].

Live-attenuated whole sporozoite approaches rely on generating large amounts of pure and aseptic live parasites for clinical grade vaccine manufacture [11]. This is a considerable bottleneck to vaccine design, testing and implementation in terms of scale, time and cost. At present sporozoites can only be isolated from infected mosquitoes via manual salivary gland dissection (SGD). As well as the obvious challenges this presents to vaccine development, difficulties with sporozoite isolation have held back general understanding of sporozoite biology when compared to blood-stages of development [16,17]. Salivary gland dissection (SGD) requires *in vivo* parasite development in the mosquito followed by manual dissection of the salivary glands 15 to 21 days post infected-blood feed. Originally described in 1964 [18] with only minor variations since [16,19–25], the dissection method involves mosquito decapitation, gland removal and homogenisation to release sporozoites. Dissection in this way is timeconsuming, taking a skilled technician over an hour to dissect 40-60 glands to a reasonable standard. With total extraction time being a critical factor for subsequent sporozoite viability [26], there is a relatively low upper limit for attaining live, infectious sporozoites. Furthermore, SGD sporozoites retain a considerable amount of mosquito-originating debris [27], known to inhibit sporozoite motility (critical for liver cell infection) [28] and act as immune modulators *in vivo* [29], potentially influencing infection progression and immune responses. Likewise, the time taken, and contamination carried over, places limits on the infectivity and development of *Plasmodium* sporozoites *in vitro* [17,22,26,28,31,32]. Rates of cell infection with *in vitro* hepatocyte (primary or hepatoma) cultures using SGD are typically less than 1% using the murine malaria model *Plasmodium berghei* [19,20,24,30] and less than 2% for human *P. falciparum* sporozoites [16,25,27,31–33]. This has been a major impediment to *in vitro* studies and screens reliant on high numbers of infection.

Several previous attempts have sought to improve throughput and purity of whole sporozoite preparation. Methods aimed at bypassing SGD have included centrifugation through glass wool [34] and compression between glass plates [35]. These alternative methods, however, have consistently yielded poor parasite purity, necessitating combination with density gradients [33,36–41]. Whilst the addition of gradients increases sporozoite yield, lengthy centrifugation times were required and final output was still contaminated with mosquito material [28,43]. Other methods trialled for sporozoite isolation have included ion exchange chromatography [42,43], and later free-flow electrophoresis (FFE) [44]. FFE is a liquid form of electrophoresis commonly used to separate organelles under native conditions based on net surface charge [45]. The poor yields or complexity of both methods, however, has limited interest in their scaled usage, despite recent developments in FFE technology in particular [46]. To date, the only scaled means for manufacture of a clinical grade vaccine has therefore relied on enlisting hundreds of skilled dissectors combined with rearing of parasites within aseptic mosquitoes.

Obtaining malaria sporozoites therefore remains a major roadblock to improving understanding of basic parasitic biology, and the scalable and reproducible production of whole sporozoite vaccines. As a response, we have developed a process that rapidly purifies both murine *P. berghei* and human *P. falciparum* sporozoites from whole mosquitoes, based on an optimized combination of homogenization, size exclusion, density and charge. Our stepwise approach facilitates the processing of hundreds of mosquitoes per hour and can be adapted to produce entirely aseptic, effectively contaminant-free sporozoites, all by a single person. The sporozoites isolated by this process show significantly improved infectivity both *in vitro* and *in vivo.* Critically, sporozoites isolated by this process offer sterile protection when given as a live-attenuated vaccine in a mouse model of infection, including, for the first time, via intra-muscular delivery. Being dissection-independent, this process will facilitate the rapid and scalable manufacture of *Plasmodium* sporozoites and could form a key enabling technology for delivery of a future effective malaria vaccine.

## RESULTS

### Rapid, dissection-independent, isolation of sporozoites from whole mosquitoes

The challenges of obtaining sporozoites for malaria research by salivary gland dissection (SGD) are a major impediment to improving understanding of the liver stages and development of effective interventions [12,24,27,30,31,51–54]. Sporozoite isolation by SGD is a low throughput and labour-intensive procedure which produces sporozoites of low purity, contaminated with mosquito-associated material. To address this need, we developed a dissection-independent stepwise process for the purification of sporozoites from whole mosquitoes (**Figure 1a**).

**Figure 1.**
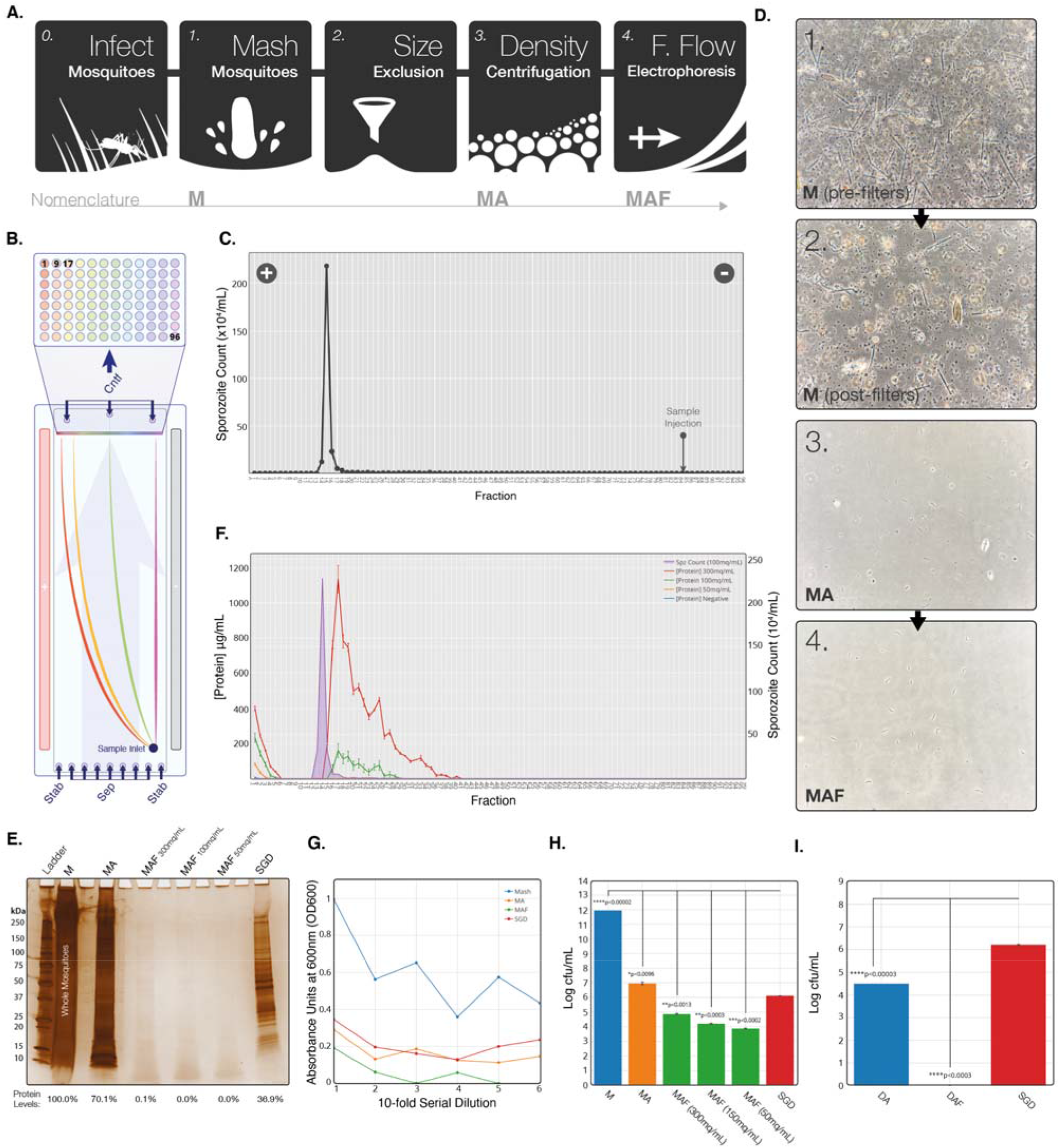
Development of a stepwise process for purification of sporozoites from whole mosquitoes. A) Schematic of key steps in the sporozoite purification process. B) Schematic representation of sample separation by cZE mode. An electrophoretic buffer is run through a chamber 0.5mm thick with a voltage applied across the flow. Sample added to the start of the chamber is carried vertically up the length of the chamber (pale blue arrow) as a voltage is applied across, separating across the horizontal length of the chamber. The outflow from the chamber is separated into 96 outlets along the horizontal length of the chamber, which drop into a 96 well plate. C) Manual sporozoite count by haemocytometer of FFE fractions from a representative MAF sporozoite separation. Point of sample injection indicated by arrow and direction of current indicated by positive and negative symbols. D) Brightfield images of each stage of purification from whole mosquito homogenate. All stages diluted to 7×10^5^ sporozoites/mL. E) Silverstain of reducing SDS-PAGE gel with <u>uninfected</u> mosquitoes (four MEQs) from each step of purification. Uninfected MAF lanes are from the same fraction as the sporozoite peak fraction identified by running infected mosquitoes at the same time. F) Protein concentration in each fraction after loading <u>uninfected</u> mosquito MA onto the FFE machine at three doses of mosquitoes (MAF). Sporozoite distribution (purple) from infected mosquitos loaded at 100mq/mL is marked to allow comparison of purification. G) End-point 16 hr serial dilution for each step of MAF purification. Absorbance of samples in TBS was measured at 600nm (OD600) 16 hr post-inoculation ay 37oC. All growth conducted at 37°C, 300rpm, using mosquitoes blood fed on <u>uninfected</u> mice 21 days prior to MAF extraction. H) Bacterial growth (samples normalised to MEQ of 200mq/mL) at different stages from <u>uninfected</u> whole mosquito (M) origin purification. Samples were loaded onto the FFE machine at three different originating mosquito doses. I) Bacterial growth (samples normalised to MEQ of 200mq/mL) at different stages from infected SGD origin purification. Experiments show the mean of two technical replicates and error bars represent SEM. All treatments compared to dissected by unpaired two tailed t-test using Bonferroni correction (H: *p<0.01, **p<0.002, ***p<0.0002, ****p<0.00002; I: *p<0.025, **p<0.005, ***p<0.0005, ****p<0.00005).

Our process consists of 4-steps, with capacity to process up to 1000 mosquitoes at a time by a single individual. Whole mosquitoes were homogenised to release sporozoites and filtered sequentially through 100μm to 10μm filters. The filtered mosquito homogenate/**M**ash (<u>M</u>) was then pre-purified by density centrifugation using **A**ccudenz (<u>MA</u>), as previously described [27], to remove larger mosquito-associated debris from sporozoites. The sporozoite layer was subsequently purified by **F**ree-flow electrophoresis (FFE) based on total net charge (<u>MAF</u>) (**Figure 1b**) using a continuous zone electrophoresis (cZE) mode (see **Methods**). Output consisted of 96 fractions with a peak sporozoite fraction, as represented by purification of *P. berghei* mCherry sporozoites assessed by light microscopy (**Figure 1c**) or fluorescent plate reader for mCherry fluorescence (**Figure S1a**).

Sporozoites produced according to this 4-step process, demonstrated reproducible separation, independent of initial sporozoite quantity. The majority of sporozoites separated into a single fraction, with a characteristic tail in the distribution that lengthened as sporozoite dose load increased (**Figure S1b-c**). This peak fraction was used for all subsequent experiments. A single FFE run, resulted in an average loss of yield of ~30% of MA input, with approximately 500-1000 mosquitoes processed by one individual in 2 hours (compared to SGD ~100 mosquitoes processed by a skilled dissector in a similar timeframe). This represents a minimum 5-10-fold increase in yield that can be run continuously or in parallel using multiple units.

Given the established potential of whole sporozoites as an effective vaccine [7] we next sought to establish the purity of MAF sporozoites compared to SGD sporozoites. Initial assessment of brightfield images clearly showed that our approach successfully removed all visible mosquito-associated debris (**Figure 1d**). To quantitively assess contaminants, samples were normalised by mosquito equivalents (MEQ); based on the number of mosquitoes (mq) homogenised and volume (units: mq/mL) as opposed to sporozoite dose, which can vary between batches. To assess the sequential reduction in protein contaminants during each step of the purification, uninfected mosquitoes were purified to determine the contributing mosquito protein contaminants. MAF samples were run at three different MEQs (300, 100, 50mq/mL) to determine an optimal purification condition. MAF purified sporozoites showed a ~100% drop in detectable protein by silver stain with <100mq/mL doses by FFE in contrast to a 63.1% reduction with SGD sporozoites (**Figure 1e**). The differences in the FFE separation profile of mosquito-associated protein at the three MEQs demonstrated that at 100mq/mL or less all detectable protein could be effectively removed from the peak sporozoite positive FFE fractions (**Figure 1f**).

Analysis of the FFE output demonstrated our ability to remove abundant mosquito proteins, such as actin, as well as enriching for sporozoite proteins in the sporozoite fraction (**Figure S1d-f**). In addition, an identical protein purification profile was obtained when using dissected salivary glands as starting homogenate rather than whole mosquitoes (referred to as **D**issected-**A**ccudenz-**F**FE; DAF) (**Figure S1d**), demonstrating the flexibility of our stepwise process for different sub-populations of sporozoite within the infected mosquito.

Given that bacterial contamination is a major problem for *in vitro* work, we next assessed the ability of our stepwise process to separate mosquito-associated bacteria. Serial dilutions of samples normalised to equal MEQ from each stage of purification were grown for 16hr at 37°C in a non-selective tryptic soya broth (TSB) medium [47] (**Figure 1g**). A marked reduction in bacterial growth, assessed by measuring OD600, was observed with MAF purified sporozoites. This was further confirmed by measuring bacteria colony forming units per mL (cfu/mL) on blood-agar plates, which showed a significant 8.1 log reduction in total bacterial load compared to a 5.9 log reduction by SGD (**Figure 1h, S1g, S2**). This translates to a 173-fold reduction in the bacterial load when compared to equivalent numbers of sporozoites obtained by SGD. Repeating the process with DAF produced sporozoites (dissected salivary glands used as input for FFE processing), demonstrated the successful removal of all detectable bacteria, effectively producing aseptic sporozoites (**Figure 1i**).

As an alternative to density gradients, we also tested whether rapid gel filtration with a Sephadex-based spin-column (5 minutes) could replace Accudenz, mirroring a method used with bovine sperm purification [48]. In parallel, we tested whether an interval zone electrophoresis (iZE) FFE method (see **Methods**) could add further improvements to our overall process (**Figure S3**). Combining Sephadex with iZE, we were able to produce completely sterile sporozoites from whole mosquitoes (**Figure S3**). Since purity was associated with an additional loss in yield (~80%), for the remainder of the development of the process (and experiments described below), MAF purification using an FFE input of 100mq/mL with the cZE method was used.

### MAF sporozoites show improved in vitro infectivity compared to those from SGD

Having developed a robust pipeline for high-throughput purification of sporozoites from whole mosquitoes with enhanced scale and purity, we next sought to assess *in vitro* infectivity. Sporozoite motility is often used as a primary indicator of sporozoite viability [24,26]. Comparisons of the motility patterns of SGD versus MAF sporozoites on a glass surface revealed no significant differences in the 2D motion patterns displayed (static, attached/waving or gliding) [49], either in terms of mean velocity or overall ratios of motion pattern (**Figure 2a-c**).

**Figure 2.**
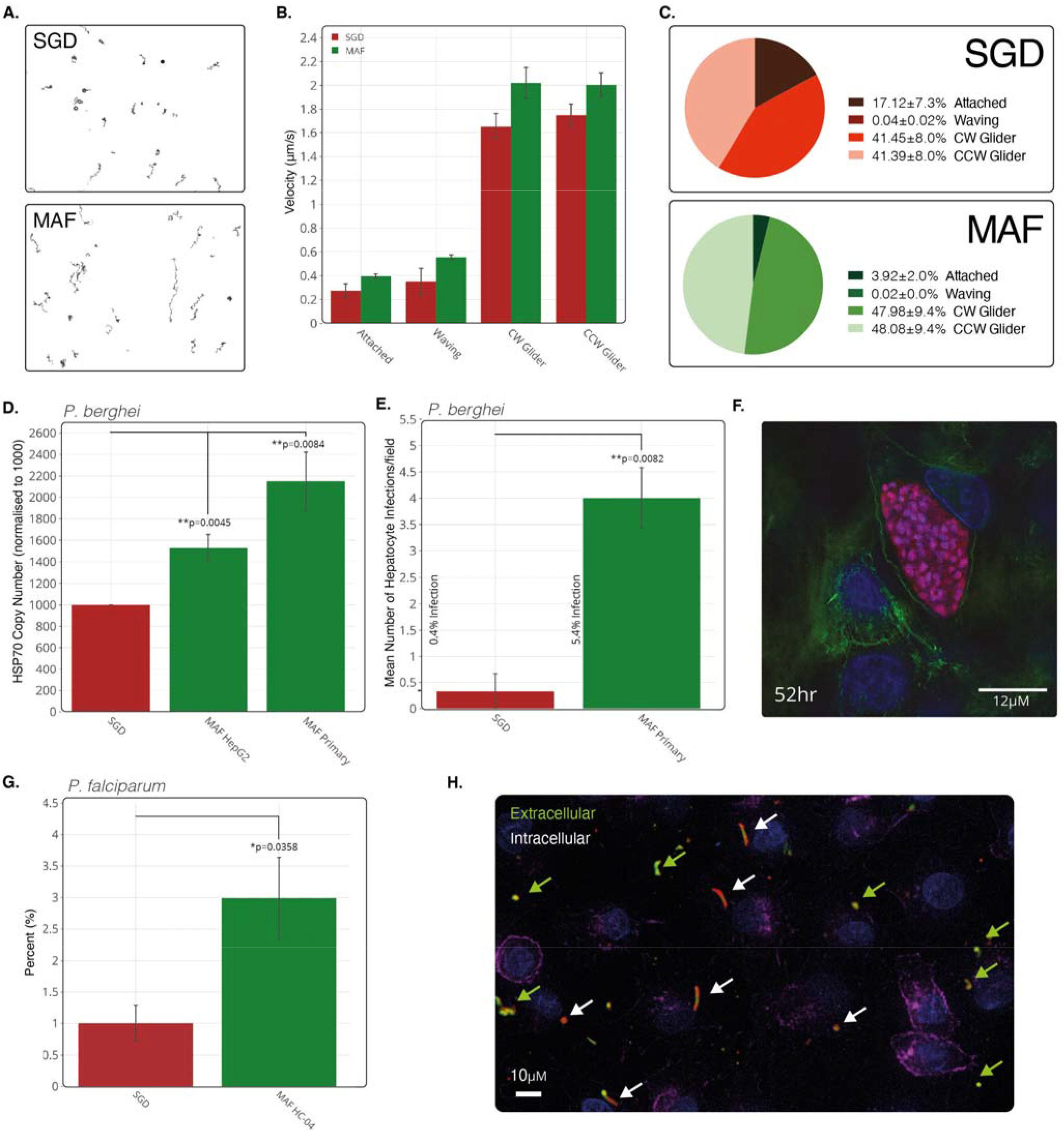
Assessment of purified sporozoite *in vitro* viability. A) Typical movement trails of MAF and SGD sporozoites over 600 frames at 2Hz. B) Sporozoite gliding motility over 600 frames at 2Hz with a sliding nine frame average during each motility state over 600 frames at 2Hz. C) Comparison of the percentage of all sporozoites in each state. Sporozoite tracking represents mean of two independent replicates and six technical replicates with groups compared using an unpaired two-tailed t-test. Bars represent means and error bars the SEM. D) Absolute RT-PCR quantification of parasite HSP70 housekeeping gene DNA copies normalised by host HSP60 gene in HepG2 and primary rat hepatocytes. Treatments normalised to 1000 *hsp70* copies for the SGD treatment. Means of three independent replicates and 6-8 technical replicates. E) Mean counts of successful hepatocyte infections in primary rat hepatocytes measured by visual identification of six fields of view over 24hr time-lapse from three independent replicates. F) Fluorescent image of late stage schizont (52hr) captured using structured illumination microscope. Blue; nuclei, green; actin, red; mCherry parasite, pink; parasite actin (anti-5H3 **[70]**). G) Means counts of successful invasions of *P. falciparum* sporozoites four hours post infection in HC-04 at a ratio of 1:5 cells to sporozoites. SGD treatment normalised to 1. Sporozoites stained for CSP to determine intracellular or extracellular location. One independent replicate with three technical replicates. H) Immunoflourescent staining of HC-04 cells with fixed four hours after infection with *P. falciparum* sporozoites and stained with anti-CSP (extracellular = green+red, intracellular = red only), DAPI for nuclear material (blue) and phalloidin for actin (purple).

Extending infectivity analysis to *in vitro* infection of hepatoma or primary hepatocytes, RT-PCR analysis of *P. berghei* copy number, 24 hours post infection (p.i.), showed a 1.5- and 2.1-fold increase in MAF sporozoite infectivity in HepG2 and primary rat hepatocytes compared to that of SGD sporozoites respectively (**Figure 2d**). More significantly, the proportion of *P. berghei* exo-erythrocytic forms (EEFs) developing in primary rat hepatocytes 24 hours p.i. was 13.5-fold increased when MAF sporozoites were used rather than their SGD counterparts (infection rate of 5.4% *versus* 0.4%) (**Figure 2e**). At 52 hours p.i., MAF sporozoites had completed maturation into late schizonts, as indicated by the presence of liver-stage merozoites (**Figure 2f**). These sporozoites also fully developed into late-stage exoerythrocytic schizonts when infected hepatocytes were extracted from rats and cultured *ex vivo* (**Figure S4**). Corroborating these results, assessment of MAF infected rat primary hepatocytes by flow cytometry, showed infection rates of 10.37% (302 out of 2808 cells) (**Figure S5**). Critically, mirroring observations in *P. berghei*, *P. falciparum-*derived MAF sporozoites were shown to have a 3-fold increased rate of invasion into HC-04 cells when compared to sporozoites isolated by SGD (**Figure 2g-h**). The absence of a robust human primary hepatocyte model for long-term *in vitro* development precluded our ability to take these to late-stage schizogony.

Overall, these results clearly show that both *P. berghei* and *P. falciparum* sporozoites purified by our method exhibit superior infectivity *in vitro*, making them preferable for *in vitro* experimentation.

### MAF sporozoites from segmented mosquitoes are more infectious than SGD in vivo

As MAF sporozoites showed enhanced infectivity *in vitro* compared to SGD counterparts, further *in vivo* studies were performed to confirm this trend. Mice were inoculated with *P. berghei* sporozoites by intravenous (i.v.) injection and infectivity determined by measuring the time to reach 1% blood stage parasitaemia (prepatent period). Mice were inoculated i.v. with escalating doses of MAF purified *P. berghei* sporozoites, demonstrating that, independent of the inoculum size, MAF sporozoites were able to develop and establish a successful blood-stage infection (**Figure 3a**).

**Figure 3.**
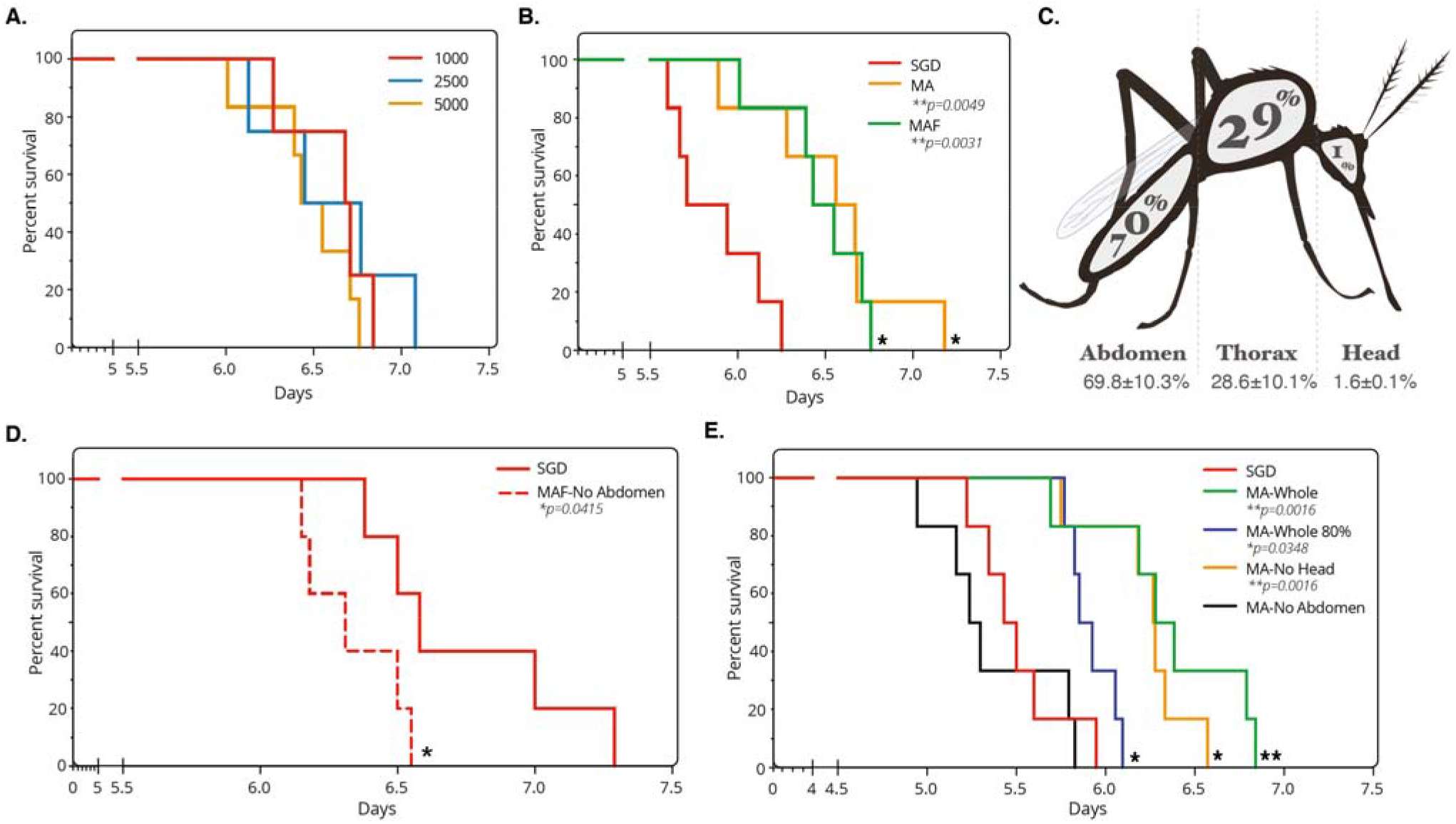
Assessment of purified sporozoite *in vivo* viability. A) Kaplan-Meier survival curve of mice challenged intravenously (i.v.) with increasing doses of sporozoites from MAF. Six mice per group. Endpoint classed as 1% parasitaemia. B) Kaplan-Meier survival curve of mice challenged i.v. with 5000 sporozoites from different purification steps. Six mice per group. Endpoint classed as 1% parasitaemia, treatments compared by Mantel-Cox statistical test. C) Sporozoite distribution of infected mosquitoes, average from 85 mosquitoes, two experimental replicates. Values show mean with SEM. D) Kaplan-Meier survival curve of mice challenged i.v. with 1000 sporozoites from MAF-No Abdomens purified (MAF from mosquitoes with abdomens removed prior to homogenisation) and SGD origin. Six mice per group. Endpoint classed as 1% parasitaemia, treatments compared by Mantel-Cox statistical test. E) Kaplan-Meier survival curve of mice challenged with 5000 sporozoites obtained by MA purification from different mosquito sources or SGD origin. Six mice per group. Death classed as 1% parasitaemia, treatments compared by Mantel-Cox statistical test.

We next compared infectivity of MAF sporozoites compared to SGD sporozoites. Initially, 5000 sporozoites obtained from either SGD, MA or MAF were given i.v. to mice with resulting blood stage parasitaemia monitored. Surprisingly, partially purified MA sporozoites and fully purified MAF sporozoites showed a modest but significant <u>delay</u> in time to 1% parasitaemia compared to SGD (0.66 days longer for MA, **p=0.0049; 0.59 days longer for MAF, **p=0.0031; Mantel-Cox Test) (**Figure 3b**). This was in contrast to *in vitro* infections which were significantly increased with MAF sporozoites. As sporozoites purified from whole mosquitoes will necessarily include a proportion of immature sporozoites that have yet to reach the salivary glands, we reasoned that this modest reduction in *in vivo* infectivity could be due to the presence of immature sporozoites in the MA and MAF inoculum. Indeed, while sporozoites obtained from the mosquito hemocoel have been shown to infect hepatocytes *in vitro* and *in vivo* [24], salivary gland sporozoites exhibit increased *in vivo* virulence compared to less mature hemocoel sporozoites [24,50]. To investigate the relative proportion of immature midgut-derived/hemocoel sporozoites in whole mosquito homogenates, mosquitoes at 21 days post infectious bloodmeal were separated into abdomen, thorax (containing the salivary glands) and head. Segmentation of the mosquito in this manner revealed that a significant number of sporozoites were indeed in the abdomen (70%) compared to the thorax (29%), which contains the salivary glands (**Figure 3c**).

We subsequently sought to determine whether subtraction of immature sporozoites from the initial inoculum could revert the delay in time to patency. Mosquito abdomens were discarded prior to the initial homogenisation step in our purification platform (MAF-No Abdomen; MAF with abdomens removed prior to homogenisation) and, as a measure of sporozoite *in vivo* infectivity, the time to 1% blood parasitaemia was monitored as before. Notably, in mice infected with MAF-No Abdomen sporozoites there was now a significant <u>reduction</u> in the time to 1% parasitaemia compared to those injected with SGD sporozoites (*p = 0.0415; Mantel-Cox Test; **Figure 3d**). This increase in infectivity was only patent when sporozoites went through all steps of the purification pipeline, as no differences were observed between mice challenged with SGD and partially purified MA-No Abdomen sporozoites (MA with abdomens removed prior to homogenisation) (**Figure 3e**). Increasing the dose of whole mosquito origin MA mosquitoes (180%, 1.8-fold increase) caused a clear reduction in the pre-patent period, indicating an increase in infectious dose. These findings correlate with *in vitro* data and demonstrate that our stepwise purification process is associated with a marked increase in sporozoite infectivity *in vivo*.

### Vaccination with irradiated MAF sporozoites confers sterile protection

Having developed a process that produces sporozoites with high purity and improved infectivity over SGD sporozoites, we next sought to assess the potential of MAF sporozoites as a radiation-attenuated sporozoite vaccine (RASv). Prior to immunisation, the effective irradiation dose was determined to be 60Gy by i.v. inoculation with varying doses of gamma irradiated sporozoites (**Figure 4a**). Mice were immunised i.v. using a three-immunisation regime of 40,000 irradiated sporozoites, two weeks apart. In parallel, cohorts of control mice were immunised with plain medium as controls. Immunisation efficacy was assessed by challenging with 5 infectious mosquito bites (**Figure 4b**) [51]. Immunisation with *P. berghei* (wildtype PbANKA) or *P. falciparum* (wildtype NF54) MAF-RASv sporozoites i.v. achieved complete protection against native *P. berghei* or chimeric *P. berghei* expressing *P. falciparum* CSP (PbANKA-PfCSP) respectively (**Figure 4c-d**). The level of protection (sterile protection) was comparable to that offered by SGD-RASv. Equivalent total IgGs measured against whole sporozoites pre-challenge were found in the serum from immunized animals irrespective of the source of *P. berghei* sporozoites (**Figure 4e**).

**Figure 4.**
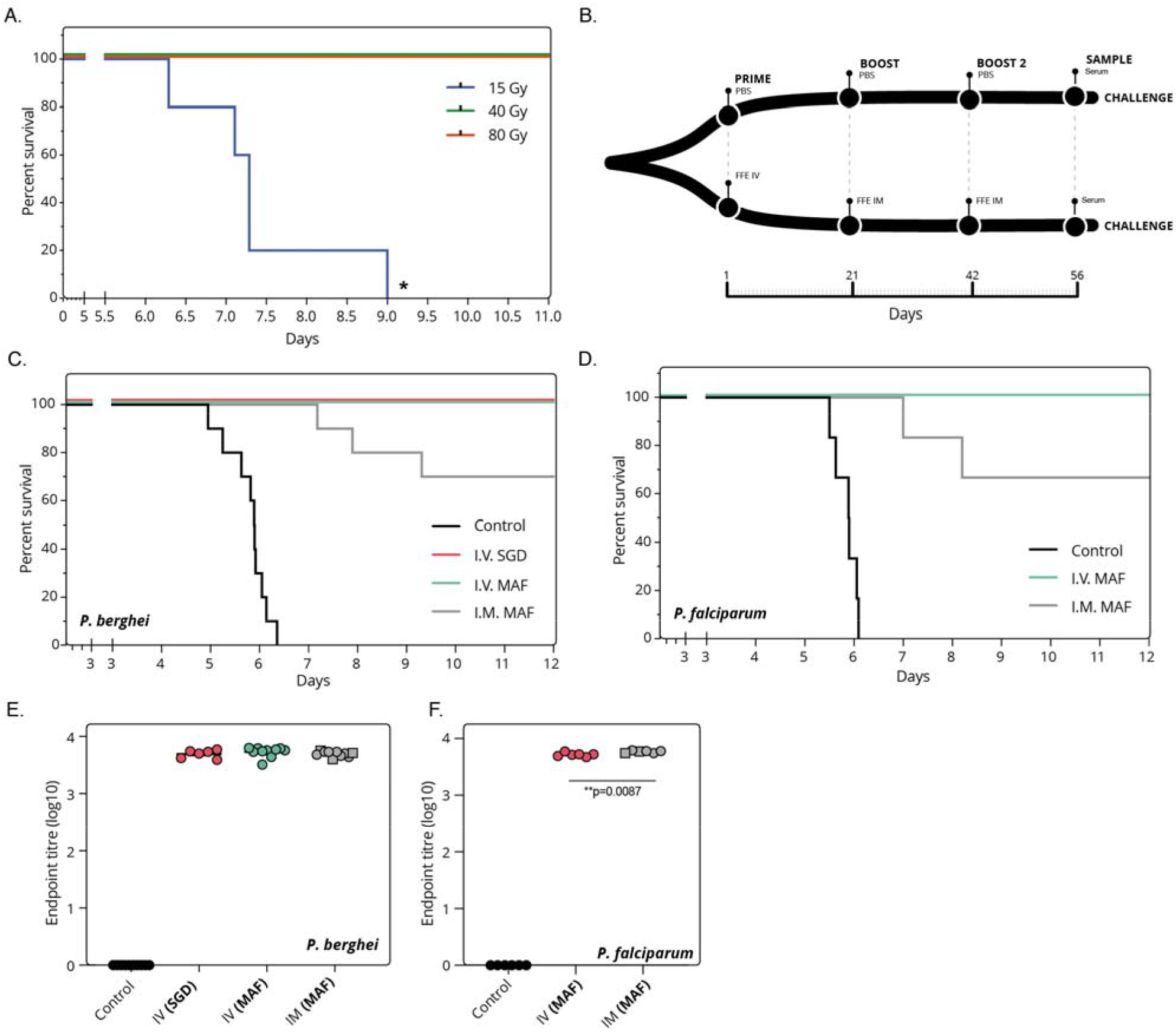
Purified sporozoites as a viable vaccine. A) Kaplan-Meier survival curve of mice challenged i.v. with 1000 *P. berghei* sporozoites from MAF-No abdomen purified and gamma irradiated. Four mice per group. Endpoint classed as 1% parasitaemia. B) Schematic of vaccination regime used. Sporozoites were either from SGD or MAF-No Abdomen origin, then gamma irradiated. C) Immunisation i.v. or i.m. of Balb/c mice with irradiated *P. berghei* sporozoites from either manual salivary gland (SGD) dissection or MAF-No Abdomen. Mice given three immunisations of 40k sporozoites, two weeks apart followed by challenge with five infectious mosquito bites. Ten mice per group. Endpoint classed as 1% parasitaemia. D) Immunisation i.v. or i.m. of Balb/c mice with irradiated *P. falciparum* sporozoites from MAF-No Abdomen. Mice given three immunisations of 40k sporozoites, two weeks apart followed by challenge with five infectious mosquito bites. Six mice per group. Endpoint classed as 1% parasitaemia. E) Total titres of IgG antibodies against *P. berghei* sporozoite lysate in mouse serum prior to challenge F) Total titres of IgG antibodies against *P. falciparum* sporozoite lysate in mouse serum prior to challenge. Squares indicate mice not protected.

Finally, considering the practical development and utilisation of a whole sporozoite vaccine and the comparatively challenging nature of i.v. administration, we sought to address whether alternative routes of immunisation such as intramuscular (i.m.) might become possible given the increased infectivity of MAF produced parasites. Previous attempts at i.m. immunisation using *P. berghei* demonstrated protection of <30% [52]. Intramuscular immunisation of 40,000 irradiated MAF sporozoites with two boosts was given with the commercial adjuvant AddaVax, a squalene-based oil-in-water nano-emulsion (InvivoGen). Remarkably, MAF sporozoites showed 70% and 67% protective efficacy for both *P. berghei* and *P. falciparum* MAF-RASv immunisation compared to controls respectively. Comparison of total IgG against sporozoites between i.v. and i.m revealed similar antibody titres for *P. berghei* immunisations (**Figure 4e**), whilst a modest increase in titre was observed with *P. falciparum* between i.m. and i.v. routes of immunization (**Figure 4f**). As the first reported demonstration of high efficacy achieved from i.m. administration for a whole-parasite malaria vaccine, these data suggest that MAF sporozoites could be an optimal starting point for scalable development of an increased efficacy and utility sporozoite vaccine for the future.

## DISCUSSION

For the first time we present an accessible, robust process for the high yield isolation of large quantities of pure malaria sporozoites without requiring manual salivary gland dissection (SGD). Sporozoites isolated by this dissection-independent process exhibited improved sterility and infectivity *in vitro* and *in vivo* and could be shown to confer sterile protection in a mouse challenge model. With demonstrated application to both murine *P. berghei* and human-infective *P. falciparum*, sporozoite production using this process represents a transformative technology that will advance a multitude of applications, not least advancing development of efficacious whole-parasite malaria vaccines.

Given the challenges of current sporozoite purification methods we set out to develop a versatile process for purification of sporozoites from whole mosquitoes; a process independent of the need for dissection. By combining bulk mosquito homogenization, Sephadex filtration (or density centrifugation) and free-flow electrophoresis separation (abbreviated to MAF), sporozoites purified using this stepwise process could be obtained 8-10 times faster than SGD and with all detectable mosquito-associated protein removed. SGD sporozoites in comparison retained ~40% of the mosquito associated protein material. In addition to the improved levels of purity, MAF sporozoites also showed a markedly improved infectivity *in vitro* for both rodent and human malaria parasites. The reduced overall time required, and consistency of production are likely to be key factors in determining *in vitro* infectivity. However, several additional factors likely account for this improvement in infectivity. When MAF and SGD sporozoites were added to primary human hepatocytes, SGD treatment was found to be associated with abnormal human cell morphology and reduced cell numbers (**Figure S6**), suggesting mosquito contaminants may be detrimental to host-cell growth, reducing overall viability of hepatocytes. In addition, we noted that MAF sporozoites had 4.3 times more cleaved CSP on their surface than SGD sporozoites (**Figure S1d**) [53,54]. Previous work has explored the importance of CSP processing on *P. berghei* sporozoite invasion. Of note, genetically altered sporozoites expressing a pre-cleaved CSP showed greater levels of *in vitro* infectivity [55]. Therefore, the acceleration of CSP processing seen following MAF purification may be a contributing factor to improved infectivity of sporozoites isolated in this way. Finally, a recent study identified the mosquito salivary protein, mosGILT as negatively modulating sporozoite motility [28]. Other mosquito-associated factors that reduce hepatocyte infectivity may be forthcoming and would be likely purified away from sporozoites using the MAF protocol, which may further contribute to improved hepatocyte infectivity. Of key interest, these factors will likely combine to help advance the relatively limited success that has been seen with *in vitro* liver stage systems that yield low numbers of infected hepatocytes following exposure to “viable” sporozoites [16,21,56–58] when compared to comparable asexual blood-stage or sexual stage high throughput *in vitro* platforms [59–62]. Further work is clearly now warranted towards this.

Although MAF sporozoites showed the same motility patterns as SGD, this was somewhat unexpected given that MAF sporozoites originate from the entire mosquito. Sporozoites originating from less mature stages, such as those in oocysts or the haemolymph, are typified by reduced gliding motility compared to mature salivary-gland resident sporozoites [24,49,50]. Indeed, we found that 70% of sporozoites in a typical day 21 post mosquito feed were abdominal in origin, conforming with previous observations [55] and indicating that a majority of MAF sporozoites would be derived from less mature oocyst or haemolymph stages. Studies on the infectiousness of sporozoite developmental stages have shown that oocyst sporozoites (i.e. within the abdominal section) are over 1000-fold less infectious than salivary gland sporozoites by intravenous challenge [50]. Thus, a significant proportion of injected sporozoites post-isolation may potentially be poorly infective, supporting a long-held belief that gliding *per se* may not be a good indicator of infectiousness. Our own infection data corroborates this, showing that use of whole mosquito homogenate was indeed associated with a significant delay in blood-stage parasitaemia when compared to production via our process but with prior abdomen removal (which reduced this delay), increasing time to patency compared to SGD (**Figure 3d**). Of note, the fact that removal of abdomens but without FFE (MA) does not advance infectivity, indicates that it is the FFE step that is critical to improving the viability of MAF sporozoites over SGD. Thus, efforts seeking to enrich specifically for highly infectious sporozoites may require prior sectioning of the mosquito to remove the abdomen but will critically still rely on the FFE step for maximal infectivity.

Sterile protection in mice is possible following administration of SGD sporozoites. Mirroring many similar studies, once irradiated, our MAF sporozoites showed full protection in both rodent and human challenge models via the i.v. immunisation route with similar antibody titres. Since it has recently been demonstrated that mosquito associated proteins are able to modify the human immune response to plasmodial sporozoites, it may be expected that such proteins could also interfere with the efficacy of a whole-sporozoite vaccines. Indeed, in agreement with this, recent work [63] showed that using Accudenz with SGD sporozoites to reduce total protein load was associated with improved pre-primed sporozoite (boost) specific CD8 T-cell responses compared to standard SGD. Thus, mosquito contaminants may cause innate immune upregulation *in vivo* with unknown, if not confounding, effects on vaccination studies. Since MAF sporozoites showed a marked reduction in mosquito-associated bacterial load, with sterility achieved using either the iZE FFE method or dissected salivary glands as homogenate, this suggests that MAF sporozoites may well outperform SGD sporozoites in immunogenicity. Our ability to gain 60-70% protection by i.m. immunization route is certainly suggestive of markedly improved immunogenicity. This is of critical importance due to difficulty in giving vaccines via the i.v. route compared to i.m. which requires a trained individual and can take up to ~5 minutes to administer in children [64]. Future work testing decreasing doses of sporozoites will be required to accurately assess comparative immunogenicity of different sporozoite sources and routes of immunization.

In conclusion, the work presented here shows the development of a complete stepwise process for purification of large numbers of highly infectious sporozoites in a rapid and scalable approach that is entirely compatible with basic biological, drug-screening and whole-parasite vaccine studies. Our process yields sporozoites at higher purity compared to those from dissected preparations alone and is associated with a marked increase in *in vitro* hepatocyte infections and enhanced *in vivo* infectivity, especially when less-mature abdominal sporozoites are removed. Sporozoites harvested by our process show markedly reduced levels of contaminants, can be produced aseptically and, critically, can be used to demonstrate high protective efficacy following both intravenous and intramuscular immunization. For basic sciences, this stepwise process will be an important step towards single cell -omic studies that require large amounts of highly pure sporozoites, free of the mosquito-associated contaminants that often limit our ability to draw conclusions from these studies [65–67]. Concurrently, application of this technology to a vaccine development pipeline, including genetically attenuated sporozoites [68,69], offers the tantalizing opportunity to develop and manufacture pure, viable, immunogenic whole-parasite sporozoite vaccines at a dramatically increased scale when compared to current methods. Production of sporozoites at scale will no doubt advance our understanding of malaria liver stage biology and help address the critical global need of an effective antimalarial vaccine.

## Acknowledgments

This work is dedicated to the memory of Shahid M. Kahn for his contributions to whole sporozoite vaccinology and for his generosity of spirit, collaborating on this work. We thank Alex Fyfe and Mark Tunnicliff for maintaining the mosquito colony, Andrew Worth for help with FACS sorting, Mark Shipman and Andreas Bruckbauer for help with microscopy, and Stephen Rothery for writing the script for automated image analysis. We also thank Annie Yang for advice on *P. falciparum in vitro* hepatocyte assays. We gratefully acknowledge InfraVec for supporting our *P. falciparum* research, in particular Roel Heutink and Geert-Jan Van for provision of infected mosquitoes required in the testing phases of this work.

## Funding

Research was directly supported by Wellcome (Investigator Award 100993/Z/13/Z, J.Baum and Career Development Fellowship 097395/Z/11/Z, AR-S), the Bill & Melinda Gates Foundation (OPP1200274, J.Baum) and an MRC research training and support grant (1240480, J. Blight). AMB thanks the MRC (New Investigator Research Grant MR/N00227X/1), PATH-MVI, Isaac Newton Trust, Wellcome Trust ISSF and University of Cambridge JRG Scheme for funding. EAl was funded by CAPES (Program Science without Borders, CSF-2361/13-2). FMA and HK thank Wellcome (OXION Strategic award 084655/Z/08/Z) for support. Microscopy work was supported through Mark Shipman in the Ludwig Institute at Oxford and the Facility for Imaging by Light Microscopy (FILM) at Imperial College London, supported by previous funding from Wellcome (Grant 104931/Z/14/Z) and the Biotechnology and Biological Sciences Research Council (BBSRC), UK (Grant BB/L015129/1). This project has received resources funded by the European Union’s Horizon 2020 research and innovation program under grant agreement No 731060 (Infravec2).

## Author Contributions

J. Blight designed and performed all experiments. Other authors helped with experimental execution, provision of reagents or analysis of data. KAS, EAt, HK, AE-T, PB, ED, and EAl helped to design and/or perform experiments. AMS, SMK and CJJ kindly provided the transgenic *P. berghei* parasite lines. FA kindly provided access to an FFE instrument. AMB, J. Baum, AR-S and AVSH helped fund, provide scientific input and design experiments. All authors contributed to manuscript preparation; J. Blight, AMB and J. Baum wrote the manuscript.

## Competing Interests

J.Blight, KAS, AR-S, AMB and J.Baum are inventors on a patent application filed internationally covering parts of this work. All other authors declare no competing interests.

## Supplementary Materials

Materials and Methods

Supplementary Figure S1-7

Supplementary Table S1-2

