## Supplementary Materials for "Dissection-independent production of a protective whole-sporozoite malaria vaccine"

**This PDF file includes:**

Materials and Methods
Figures. S1 to S7
Tables S1 to S2

**Materials and Methods:**

<u>Mosquito Maintenance:</u> *Anopheles. stephensi* mosquitos used for experiments were raised at 28˚C, 70% relative humidity with a 12hr light cycle. Larvae were fed with fish pellets and adults maintained on 10% fructose. Reared by Alex Fyfe and Mark Tunnicliff.

<u>*P. berghei* Maintenance and Infection:</u> Two transgenic *P. berghei* ANKA lines were used in this study that express either mCherry or GFP under control of the *uis4* promoter. This promoter drives transgene expression specifically in sporozoites and liver stages. The transgene expression cassettes of both lines has been introduced into the neutral p230p gene locus by the method of GIMO transfection [71]. The generation and characterization of the mCherry-expressing line mCherry@Pbuis4_230p (line 2204) has been described previously [72]. The generation and characterization of the GFP-expressing line GFP @Pbuis4_230p (line 2227) was generated as follows: The *P. berghei* ANKA line GIMO parent line 1596cl1 [71] was used for transfection with a construct (pL1962) which targets the neutral *p230p* locus (PBANKA_030600) and inserts GFP::Luciferase expression cassette, thereby removing the selectable marker (SM) consisting of human dihydrofolate reductase and the yeast cytosine deaminase and uridyl phosphoribosyl transferase (h*dhfr::yfcu*), according to the GIMO (gene insertion/marker out) transfection technique, which has previously been described [71]. The transfection vector, which lacks a drug selectable marker cassette, was obtained using the using the standard GIMO DNA construct pL0043 [71]. The expression cassette contained the GFP::Luciferase flanked by the 5′ and 3′ promoter and transcription terminator sequences of *P. berghei uis4* gene (PBANKA_0501200), which were amplified from *P. berghei* ANKA wild-type (WT) genomic DNA. The regulatory sequences of *uis4* gene were chosen to express GFP::Luciferase in sporozoites and liver stages [73,74]. Sequences of primers used for pL1962 construct generation are listed in **Table S1.** Transfection (exp. 2227) of 1596cl1 was performed using standard transfection methods [75] and negative selection was applied by treating mice with the 5-fluorocytosine (5-FC) in drinking water as described for GIMO transfection

[71]. The selected parasites were cloned by limiting dilution in mice and line 2227cl6 was further characterized for correct integration of GFP::Luciferase expression cassette into the *p230p* locus by diagnostic PCR and Southern analysis of pulsed-field gel electrophoresis-separated chromosomes [75]. Sequences of primers used for PCR genotyping are listed in **Table S2.** The selected parasite 2227cl6, named GFP::Luc@Pbuis4_230p, contains the fusion gene *gfp-luciferase* under the control of the *uis4* regulatory sequences integrated into the neutral p230p locus and is SM free (**see Figure S7**). For vaccination studies mice were immunised with either the PbANKA 2.34 (wildtype) or NF54 (wildtype). Subsequent challenge was with either PbANKA 2.34 or PbANAKA 2.34 transgenic for *P. falciparum* CSP (PbANKA-PfCSP chimeric[76]).

For infection of mice, cryopreserved parasitized RBC's (day five) were thawed and injected into naïve Balb/c mice by the intraperitoneal (i.p.) route and *An. stephensi* mosquitos allowed to feed on anesthetised mice with 1-2% blood-stage parasitaemia. 7-10 days later these mosquitoes were allowed to take an additional bloodmeal on naïve Balb/c mice to increase sporozoite yields. Blood-fed mosquitos were maintained at 19°C at 70% relative humidity for 19-22 days before sporozoites were extracted.

<u>*P. falciparum Maintenance and Infection:*</u> The wildtype NF54 *P. falciparum* strain was used in this study and cultured *in vitro* and gametocytes induced as per Delves *et al* [77]. Briefly, asexual cultures were grown in RPMI 1460, supplemented with 25 mM HEPES (Life Technologies), 50 µg L-1 hypoxanthine (Sigma) and 10% A+ human serum (Interstate Blood-Bank). Gametocyte cultures were grown in RPMI 1640 supplemented with 25 mM HEPES (Life Technologies), 50 µg L-1 hypoxanthine (Sigma), 2 g L-1 sodium bicarbonate (Sigma), 5% A+ human serum (Interstate Blood-Bank) and 0.5% AlbuMAX II (Life Technologies). For standard membrane feeding assays, 15-17 day-old gametocyte cultures were diluted in fresh RBCs and human serum at 50% haematocrit and used to feed female

overnight starved mosquitoes. Mosquitoes were maintained for 16-18 days before sporozoites were extracted.

*Manual Salivary Gland Dissection:* Mosquitoes were sedated on ice for 10 min, then placed on a glass slide with 100µL complete Schneider's *Drosophila* medium (1% FBS, 4°C, $NaHCO_3$ free, Pan-Biotech) and whole salivary glands removed by gentle separation of the head using micro-forceps. Both sets of glands were gently cleaned to remove other tissues then placed into a glass dounce tissue grinder on ice using 2µL fresh medium. The glass slide was cleaned between each dissection. Each dissection took approximately (45-90 sec) and was carried out for no more than 2-3 hr maximum to reduce loss of infectivity. To release sporozoites the salivary glands were homogenised with three gentle but firm grinds using the pestle. The sample was transferred to protein lo-bind tubes (Eppendorf) used to prevent loss of sporozoites by adhesion to plastic-ware and mixed well before a sample was added to a haemocytometer and the average of four 16 square fields counted. Sample was diluted if too concentrated to accurately count.

*Homogenisation and Accudenz Gradient Purification/Sephadex:* Mosquitoes sedated on ice were placed in a Petri dish with 2mL (per 400 mosquitoes) complete Schneider's *Drosophila* media and gently homogenised with the end of a 10mL syringe barrel for 30-60 sec or using a gentleMACS homogeniser (miltenyl biotech). Liquid was removed and passed through a 100µM cell strainer in a 50mL centrifuge tube. A further 1.5mL media was added to the petri dish, gently homogenised and passed through the 100µM filter. This was repeated twice more but with more vigorous grinding. Finally, the filter was washed with 1mL media. The filtrate was subsequently passed through a 70, 40 and 20µM filter and each washed with 1mL media (M). All steps were carried out on ice. On some occasions, the head and/or abdomens were removed before homogenisation. 1mL homogenate was loaded onto a 3mL Accudenz cushion (4°C) in a 15mL centrifuge tube and centrifuged (2500 xg, 4°C) as per reference [27].

Subsequently, 400μL was taken from the sporozoite enriched boundary (at the 3mL mark). 1mL aliquots of Accudenz sporozoites were put into 2mL protein lo-bind tubes (Eppendorf), made up to 2mL with complete Schneider's media and centrifuged (12,000 xg, 4$^{o}$C, 3 min). The resultant pellet was re-suspended in complete Schneider's media (MA). If the sample was to be used for free-flow electrophoresis (FFE) it was re-suspended to a mosquito equivalent (ME) of 200mq/mL (mosquitoes/mL; unless stated otherwise), which was based on the original number of whole mosquitoes homogenised and the final volume this was in. The resultant sporozoite suspension was in most cases was further purified by FFE.

Alternatively, SG-15 Sephadex medium was prepared in 3% sodium citrate at a 1:1 v/v ratio overnight. Subsequently a PD10 column was packed with 5cm depth Sephadex and homogenate applied. This was centrifuged for 1-5 minutes at 500-1500rpm. Eluted samples were subsequently applied to FFE.

*Free-Flow Electrophoresis Purification:* Prior to sporozoite extraction the FFE machine (FFE Service GmbH) was setup for continuous ZE (cZE) using a 0.5mm ZE spacer. A separation buffer of 10mM triethanolamine (TEA), 10mM glacial acetic acid (HAc) and 250mM sucrose was used with a stabilisation buffer of 100mM TEA, 100mM HAc and 250mM sucrose injected into the separation chamber at 150/300mL/hr. Electrodes were kept in 100mM TEA, 100mM HAc and 250mM sucrose with a voltage of 900/750V and current and power limit of 150mA and 150W respectively. MA sample was mixed 1:1 with separation buffer (now at 100mq/mL) and injected into the separation chamber at the cathode end at a rate of 1600μL/hr and fractions collected 14 min after injection started and stopped 14 min after sample finished. Fractions were collected at 4˚C in 2mL protein lo-bind deepwell plates (Eppendorf) containing 400uL complete Schneider's medium. The peak sporozoite fraction(s) was identified by a haemocytometer and centrifuged in 2mL protein lo-bind tubes (max, 4$^{o}$C, 3 min) and the pellet re-suspended in 100-500µL complete Schneider's media (MAF). To compare purification stages all samples were re-suspended to the same ME. FFE ME dose was calculated based on the volume collected in the peak fraction. Alternatively, the FFE machine was setup for interval zone electrophoresis (iZE) using a 0.2mm spacer with identical separation buffers. Settings were 1200V, 150mA, 120W with injection at 2000 uL/hr.

*Hepatocyte Culture:* Tissue culture plates were pre-coated overnight or using plasma-treatment with a 0.1M bicarbonate buffer (pH9.4) [78] of collagen I, collagen IV, fibronectin and laminin (Sigma-Aldrich; 1µg/cm$^2$). Human HepG2 hepatoma cell lines were maintained in complete DMEM (10% FBS, 1% penicillin/streptomycin, 5% L-glutamine; Sigma-Aldrich) at 37$^o$C with 5% $CO_2$. A confluent monolayer was maintained using a 18G syringe needle. HC-04 cells were maintained in complete medium (DMEM supplemented with F12, 10% FBS, 1% penicillin/streptomycin, 5% L-glutamine; Sigma-Aldrich). To obtain primary hepatocytes male Wistar rats [Crl:CD(SD), strain 001] were anesthetised and a 21G cannula was inserted into the hepatic portal vein and secured/sealed using tissue adhesive (3M). Liver perfusion medium (Thermo Sci; 37$^o$C) was pumped through the cannula at 10mL/min using a peristaltic pump and once the liver started to lighten (within 30 sec) the speed was adjusted to 20mL/min. Subsequently the inferior vena cava was cut and over the next 5 min blocked 2-3 times and the pump increased to 40mL/min. Following successful perfusion, the media was exchanged for liver digest medium (Thermo Sci; 37$^o$C) and the same blocking procedure carried out for 8 min. The liver was subsequently transferred quickly to 4$^o$C complete DMEM on ice, the liver disrupted (using gentleMACS homogeniser) and the passed through 100µM cell strainers. The cell suspension was washed twice (50 xg, 5 min, 4$^o$C) with a final re-suspension into 19mL complete DMEM and 20mL sterile isotonic percoll (SIP; 90% percoll, 10% 10xPBS) and centrifuged (1.06g/mL, 100 xg, 10 min, 4oC) to remove debris and dead cells (percoll purification modified from reference [79]). The pellet was washed in complete DMEM and used to seed plates. Importantly the plates were not moved for 30 min to allow the cells to adhere evenly across the plate. They were then transferred to an incubator (37°C, 5% $CO_2$) for 1-2 hr before medium was exchanged with serum-free hepatocyte growth medium (Promocell) which was exchanged every 12-15 hr.

<u>Sporozoite Motility Assessment:</u> Sporozoites were added to 37°C complete DMEM and centrifuged (1,500 rpm, 4 min) in glass bottom tissue culture plates to sediment sporozoites. Fluorescent images were captured at 2Hz for 600 frames at 20x magnification. Motility was assessed using the ToAST ImageJ plugin [49].

<u>Murine Sporozoite Challenge:</u> *P. berghei* sporozoites were extracted from infected mosquitoes using one of the described methods and diluted in complete Schneider's *Drosophila* medium (1% FBS, 4°C). Mice were placed in a 37°C heat-box for 10 min prior to injection of 50µL intravenously (i.v.) into either lateral tail vein of restrained mice. From day 4-5 parasitaemia was monitored by thin-blood film until three days of positive smears were obtained, mice were then sacrificed. Time to 1% parasitaemia was then calculated by linear regression. If parasites were not detected by day 14 the mice were sacrificed.

<u>Murine vaccination:</u> Prior to immunisations sporozoites were diluted in Schneider's *Drosophila* media to $80x10^4$/mL and irradiated using a Cs-137 gamma irradiation source with a Gammacell 3000. For intravenous immunisations sporozoites were diluted to $40x10^4$ sporozoites/mL in Schneider's *Drosophila* medium and 100µL injected per mouse. For intramuscular immunisations diluted to $40x10^4$ sporozoites/ml of Schneider's media and mixed with equal volumes of AddaVax adjuvant (InvivoGen). Balb/c mice were immunized intramuscular with 50ul per site sporozoite/adjuvant mixture at 0, 3 and 5 weeks. The mice were challenged with five PbANKA 2.34 or PbANKA-PfCSP chimeric infected mosquito bites one week after booster immunization by allowing the mosquitoes to feed on the abdomen of each mouse for 15min. The salivary glands from all blood-fed mosquitoes have been

dissected after the bites to confirm the presence of infective sporozoites. Since day 4 post challenge, the immunized mice were checked daily for the presence of *P. berghei* blood stages by microscopic examination of Giemsa stained thin smears of tail blood. The mouse was classified as negative for infection when no blood-stages of parasite were present on day 14 after challenge.

*Enzyme-linked immunosorbent assays (ELISAs):* Sporozoite lysate was prepared by pelleting MAF purified sporozoites and flash freezing, before diluting in PBS and using to coat 96 well immunosorbent plates (NUNC MaxiSorp) overnight (1,500 sporozoites per well). Subsequently liquid was removed and wells allowed to air dry before blocking with 1% BSA in PBS. Wells were subsequently incubated with mouse serum with starting dilutions of 1:50 or 1:100 in 0.01% Tween-PBS. Anti-mouse IgG secondary antibody conjugated to alkaline phosphatase (AP) (Sigma Aldrich) was added after 5 washes in 0.01% Tween-PBS. AP was quantified after 5 washes in 0.01% Tween-PBS using 4-Nitrophenyl phosphate disodium salt hexahydrate (Sigma Aldrich) measured at 405nm absorbance.

*In vitro hepatocyte infection:* *In vitro P. berghei* sporozoite infections were carried out on HepG2 or primary hepatocytes 24 hr after plating. Sporozoites in 4$^{o}$C complete Schneider's *Drosophila* media were diluted in pre-warmed (37$^{o}$C) complete DMEM (for HepG2; Sigma-Aldrich) or primary hepatocyte medium (for primary hepatocytes; Promocell) to achieve a desired ratio of sporozoite to hepatocyte (usually 1:1 or 1:2) and the culture media was exchanged with the sporozoite media. Cell cultures were carefully returned to the incubator to prevent swirling and an uneven distribution of sporozoites. Media was then no-longer exchanged for the remainder of the experiment. For *P. falciparum* infections, HC-04 cells (media composition according to Yang *et al* [80]) were plated on 96 well plates coated with a 1μg/cm$^2$ mix of collagen, fibronectin and laminin (see above). Sporozoites were added to cells at a ratio of 5:1. The plates were immediately spun down for 5 min at 3000 rpm, before being returned to the incubator. After 4 hours, the cells were washed once with PBS and fixed with 4% PFA.

*Ex vivo hepatocyte challenge:* Rats were i.v. challenged with 30 million GFP transgenic sporozoites and 14 hr later hepatocytes extracted by liver perfusion (above). Infected (GFP positive) hepatocytes were sorted (MoFlo) and plated for up to 30 hr.

*Bacterial Contaminant Quantification:* To assess the sterility of each purification step, tryptic soya broth (TSB; Oxoid) was inoculated with samples normalised by MEQ and absorbance at 600nm measured after 16hr incubation at 37$^{o}$C. Alternatively, samples normalised by MEQ were serially diluted in PBS and spread on blood-agar plates incubated overnight at 37$^{o}$C. Negative growth was confirmed by a further 24hr incubation.

*Protein Purity Quantification:* For western blotting, sample was lysed using RIPA buffer with protease inhibitor cocktail (Sigma-Aldrich), protein concentration normalised using Pierce BCA protein assay kit (Thermo Scientific) and sample loaded onto a 12% TGX SDS-PAGE gel using reducing Laemmli buffer and transferred by semi-dry transfer onto a PVDF membrane (Bio-rad Laboratories). *P. berghei* CSP protein was probed using the 3D11 monoclonal [53] and detected using HRP chemiluminescence. Total protein concentration of purified mosquito sample was assessed in SDS-PAGE gels using Pierce silver stain kit (Thermo Scientific) or in solution using a Pierce BCA protein assay kit (Thermo Scientific). Dot blots were conducted on FFE fractions by loading 200uL of each fraction onto a multiscreen-IP plate (0.45μM, Millipore) pre-activated with methanol and incubated overnight (4$^{o}$C) before probing and detection of anti-mosquito actin (Sigma, A2066) similar to western blotting using HRP and ECL. Alternatively, protein contaminants were assessed using liquid chromatography tandem-mass spectrometry (LC-MS/MS) with prior sample preparation in 6M urea, 100mM tris (pH7.8), 5mM dithiothreitol, 20mM iodoacetamide with subsequent trypsin digestion overnight and desalting. Mass spectrometry output data was analysed using the Mascot algorithm (V2.4) and UniProt database. All media used for protein assessment was protein free. Equivalent volumes were injected onto FFE and collected for each treatment and total protein in each fraction quantified.

*Flow Cytometry:* Flow cytometry was carried out using an LSRII (Becton Dickson). Hepatocytes were washed three times in 1x PBS and removed by gentle cell scraping. Hepatocytes were gated for single cell using FSC-H versus FSC-A and mCherry-*P. berghei* infected cells detected in the PE-Texas Red channel by comparing to APC channel auto fluorescence. Uninfected hepatocytes were run as controls. GFP-expressing *P.berghei* infected primary hepatocytes were sorted using a MoFlo cytometer (Beckman) gated for GFP positive single cells.

*Immunofluorescent staining:* Cells were fixed with 4% paraformaldehyde and permeabilised using 1% Triton-X100. Prior to antibody incubation cells were blocked with 1% BSA and then probed with primary and then secondary antibodies in 1% BSA for 1-2 hr. Nuclear staining was carried using DAPI. For CSP in/out staining fixed cells were probed with CSP antibody before and after permeabilization.

*Fluorescent Microscopy:* Imaging of mCherry fluorescent *P. berghei* infected primary hepatocytes was carried out using 1.5mm glass bottom dishes/plates (Mattek) on a widefield fluorescent microscope with LED fluorescence light source at 2Hz using the Metamorph software package (Ludwig Institute, Oxford). Infections numbers determined by manual counting using the mCherry channel. Late stage schizonts were captured using structured illumination microscopy with a Zeiss, Elyra (Imperial College London, FILM facility). *P. falciparum*-infected cells were imaged on a Nikon Eclipse Ti (Imperial College London, FILM facility). Image processing and analysis was automated by running a custom macro in Fiji. Quantification of intracellular versus extracellular parasites was carried by determining the area fraction (as a %) of CSP in and outside of hepatocyte. Parasites were classed as inside cells if the area fraction of CSP outside and inside was <10% and >80% respectively. HC-04 were numbers were quantified using nuclear count. Cell infection was calculated as the % ratio of intracellular parasites to HC-04 nuclei.

<u>*Quantitative PCR:*</u> DNA was extracted from cultures using phenol-chloroform-isopropanol precipitation and re-suspended in molecular grade water. Nucleic acid concentration was determined using a Qubit fluorometer (Thermo Scientific). Quantification of *P. berghei* hepatocyte infection density based on absolute genome copies was determined using a standard curve plasmid containing a 271bp fragment from murine heat shock protein (HSP) 60 (Ensembl: ENSMUST00000027123) housekeeping gene and a 176bp fragment from *P. berghei* HSP70 gene (PBANKA_071190). 100ng of DNA template was amplified using SsoAdvanced Universal SYBR green supermix (Bio-Rad Laboratories) run on a CFX Connect RT-PCR machine (Bio-Rad Laboratories) as per manufacturers standard protocol and parasite genome numbers determined using linear fit normalised to HSP60 housekeeping (HSP60 HepG2 F: GACCAAAGACGATGCCATGC, R: GCACAGCCACTCCATCTGAA; HSP60 Rat F: TGGAGAGGTCATCGTCACCA, R: CACAGCTACTCCATCTGAGAGT; HSP70 *P.berghei* F: AGGAATGCCAGGAGGAATGC, R: AGTTGGTCCACTTCCAGCTG).

<u>*Animal Research:*</u> All animal work in this thesis was carried out according to the Animals (Scientific Procedures) Act 1986 Amendment Regulations 2012 (SI 2012/3039) with approval from the University of Oxford and Imperial College London Ethical Review Committee (PPL 30/2889 Oxford, 70/8788 and PDA3EBA4A Imperial). The Office of Laboratory Animal Welfare Assurance for Imperial College covers all Public Health Service supported activities involving live vertebrates in the US (no. A5634-01). Rats and mice were kept in individually ventilated cages.

*Statistical Analysis:* Data was assessed for normality and equality of variance and used to determine the suitable statistical test as per John Tukey's exploratory data analysis method [81]. Parametric data was assessed using a t-test and non-parametrically using a Mann-Whitney U test for single treatment comparisons. Multiple treatments were compared using a t-test with Bonferroni correction. Kaplan-Meier curves were compared using the Mantel-Cox test.

**Supplementary Figures and Tables:**

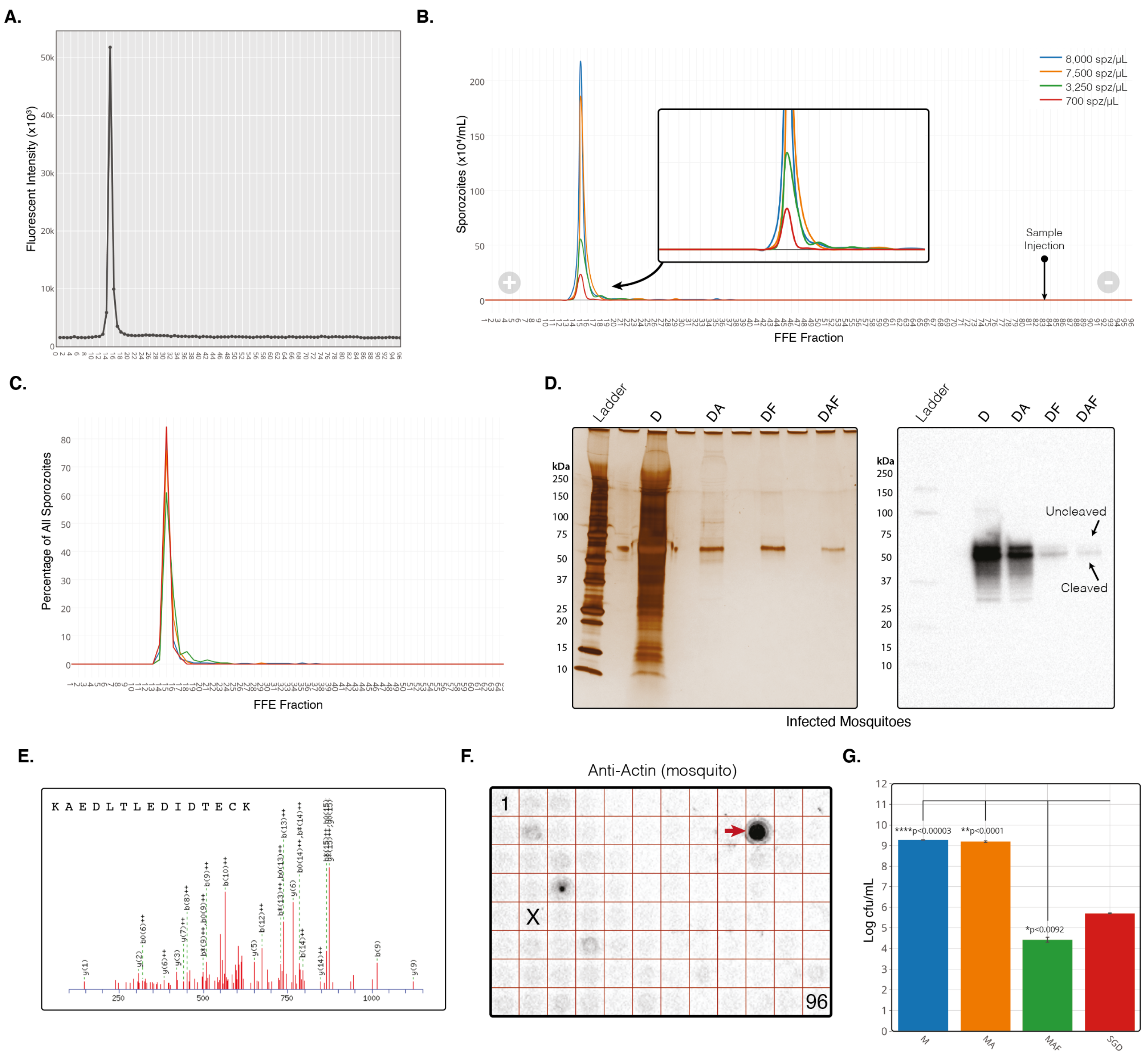


**Figure S1 | The MAF Purification platform additional data.** A) Fluorescent plate read at 610 emission of FFE fractions from a representative MAF sporozoite separation. B) Parasite distribution into FFE fractions when loaded at four different sporozoite doses (spz/mL). Quantification by haemocytometer count. C) Sporozoite distribution based on total percent of sporozoites per fraction for each sporozoite dose. D) Left; Silverstain of infected mosquitoes from each step of purification when using dissected salivary glands for homogenisation instead of total mosquitoes. Right; western blot against *P. berghei* CSP from the same samples as the silver stain (left). MAF samples in D) where injected into the FFE machine at 100mq/mL. All silver stains from reducing SDS-PAGE's with samples normalised by MEQ with 4 MEQs loaded onto each lane. Following MAF purification the ratio of uncleaved:cleaved was 1:7.8, compared to 1:1.8 for SGD sporozoites (assessed by densitometry). Cleaved and uncleaved bands indicated by arrows, with expected sizes of 45 and 55kDa respectively. E)

Liquid chromatography tandem mass spectrometry (LC-MS/MS) analysis of each stage of purification. LC-MS/MS raw data searched against the Uniprot-Swissprot database using the MASCOT search algorithm identified *P. berghei* CSP protein exclusively with three peptides in the fraction purified by MAF only. Identification of tryptic peptide 297-312 from *P. berghei* CSP protein (Swissprot ID: P06915) with MASCOT score of 37 by mass spectrometry analysis. All samples normalised to 200mq/mL. F) Dotblot of all 96 FFE fractions in a plate layout against mosquito actin protein (Sigma Aldrich, A2066). Actin positive fraction indicated by arrow. Sporozoite peak fraction indicated by X Fraction 1 and 96 indicated. G) Bacterial growth at different stages from infected whole mosquito (M) origin purification.

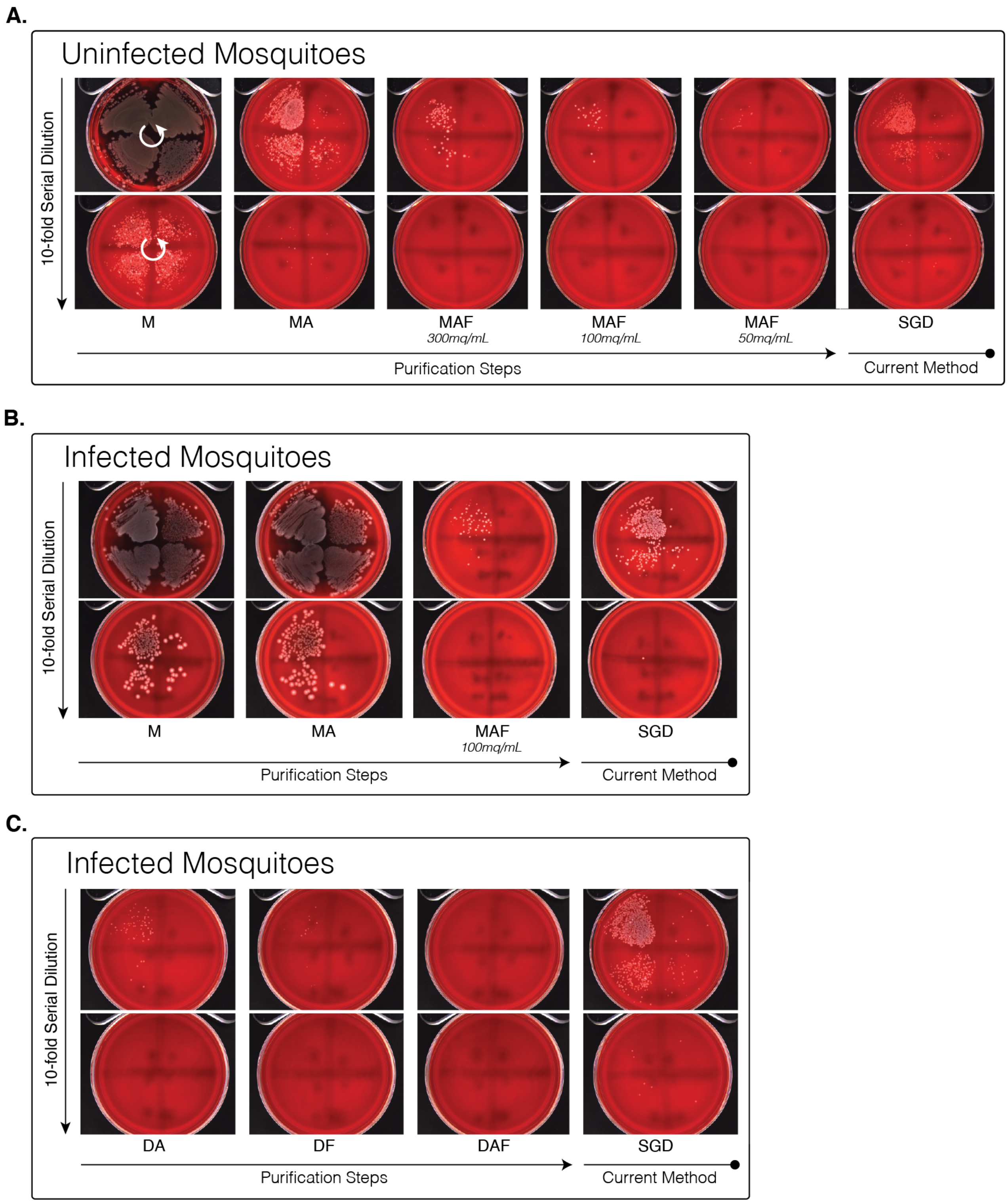


**Figure S2 | Blood plate agar growth of sporozoite purification steps.** A) Bacterial growth at different steps of purification from uninfected whole mosquito homogenate. Samples were loaded onto the FFE machine at three different MEQs (300, 100, 50mq/mL). B) Bacterial growth at different purification steps from infected whole mosquito homogenate. C) Bacterial growth at different stages from infected dissected salivary gland homogenate. All samples spread onto blood agar plates in eight, 10-fold serial dilutions running anti-clockwise, with plates separated into 4 quadrants and dilutions starting in the top right quadrant (indicated by white arrows).

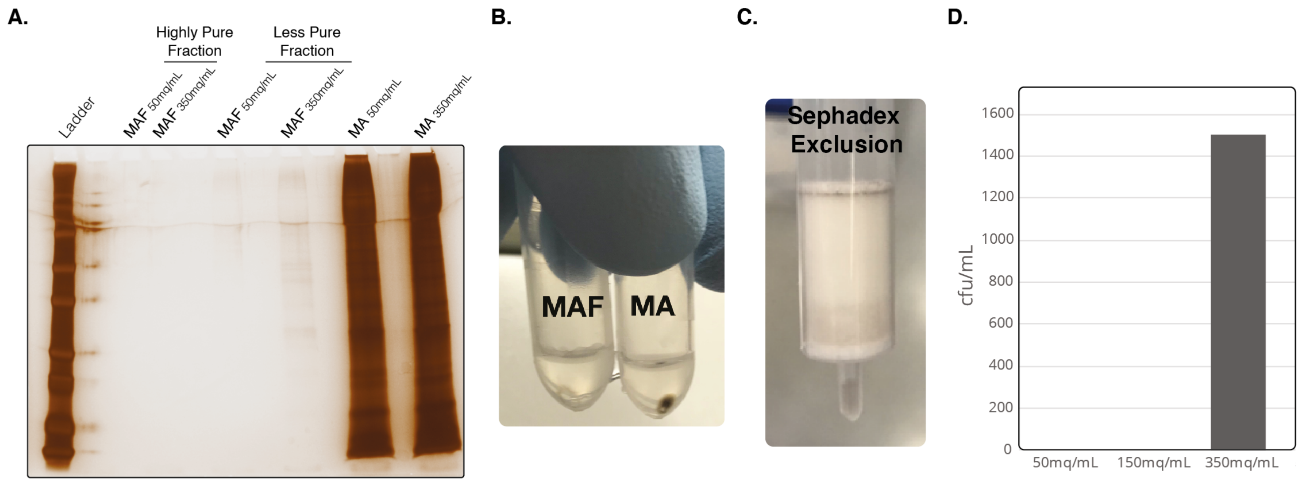


**Figure S3 | MAF Purification using Sephadex and iZE methods.** A) Silver stain of each stage of purification illustrating the high and low purity fractions. All samples run at identical MEQs. B) Images of pellets from MA and MAF purification. C) Images of a Sephadex column after eluting sample. D) Bacterial cfu/mL of samples injected into the FFE (iZE) at three concentrations of MEQs.

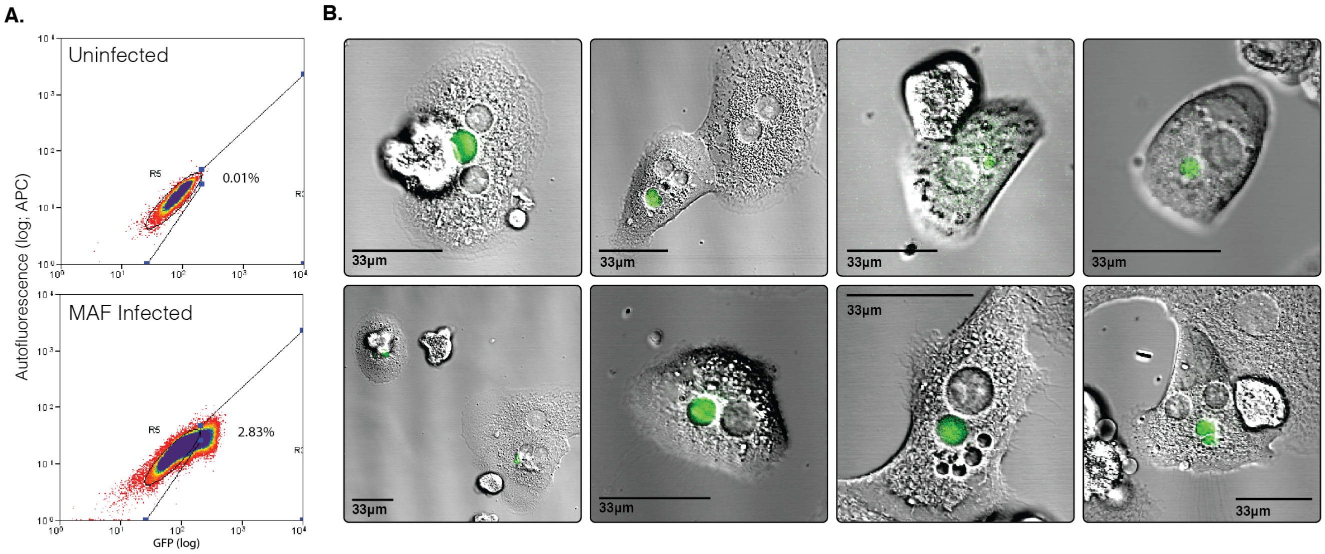


**Figure S4 | *Ex-vivo* development of MAF purified *P. berghei* sporozoites in rat primary hepatocytes.** A) The ability of MAF sporozoites to infect hepatocytes *in vivo* but develop *ex vivo* was investigated. Hepatocytes were extracted by perfusion of livers that were collected from rats 14 hr after intravenous injection of sporozoites into rats. These rats were infected with a total of $3x10^7$ GFP-expressing *P. berghei* sporozoites purified by MAF (whole mosquitoes) from 400 mosquitoes. Infected hepatocytes from these rats were collected by flow-sorting and subsequently plated and incubated for a period of up to 30 hr. Flow sorting identified 2.83% GFP-positive cells in the extracted, perfused liver cell population. B) Fluorescent images of GFP positive cells collected by flow sorting 24 hours after plating.

A.

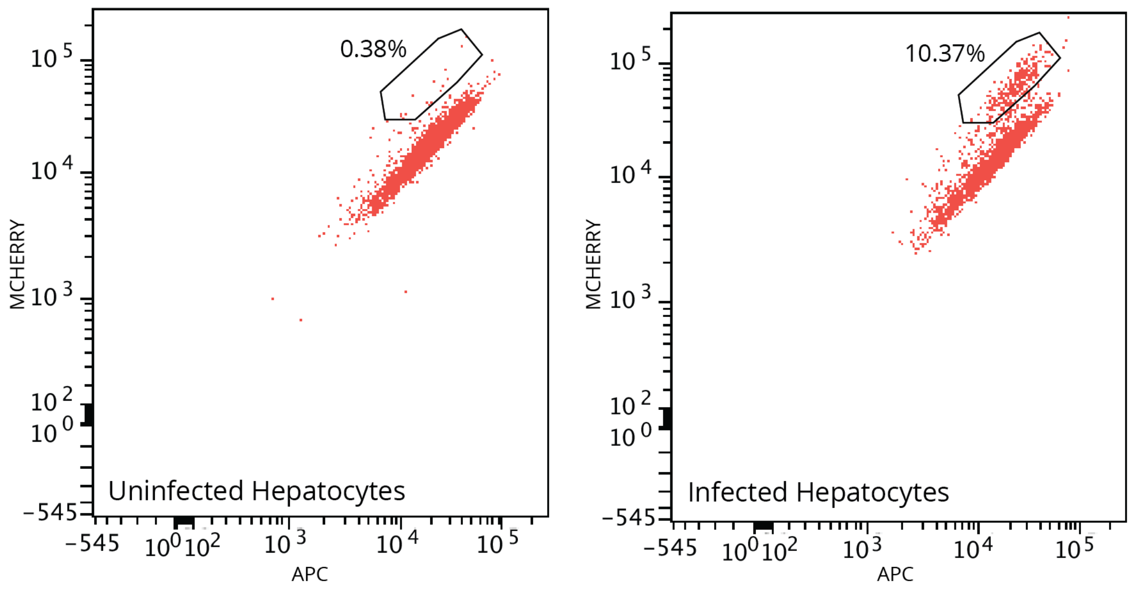


**Figure S5 | MAF *P. berghei* flow cytometry.** A) Flow cytometry quantification of mCherry expressing transgenic *P. berghei* infected primary rat hepatocytes 36 hr post addition of sporozoites (total 2808 cells in treatment, 3906 cells in control). Data representative of three technical replicates

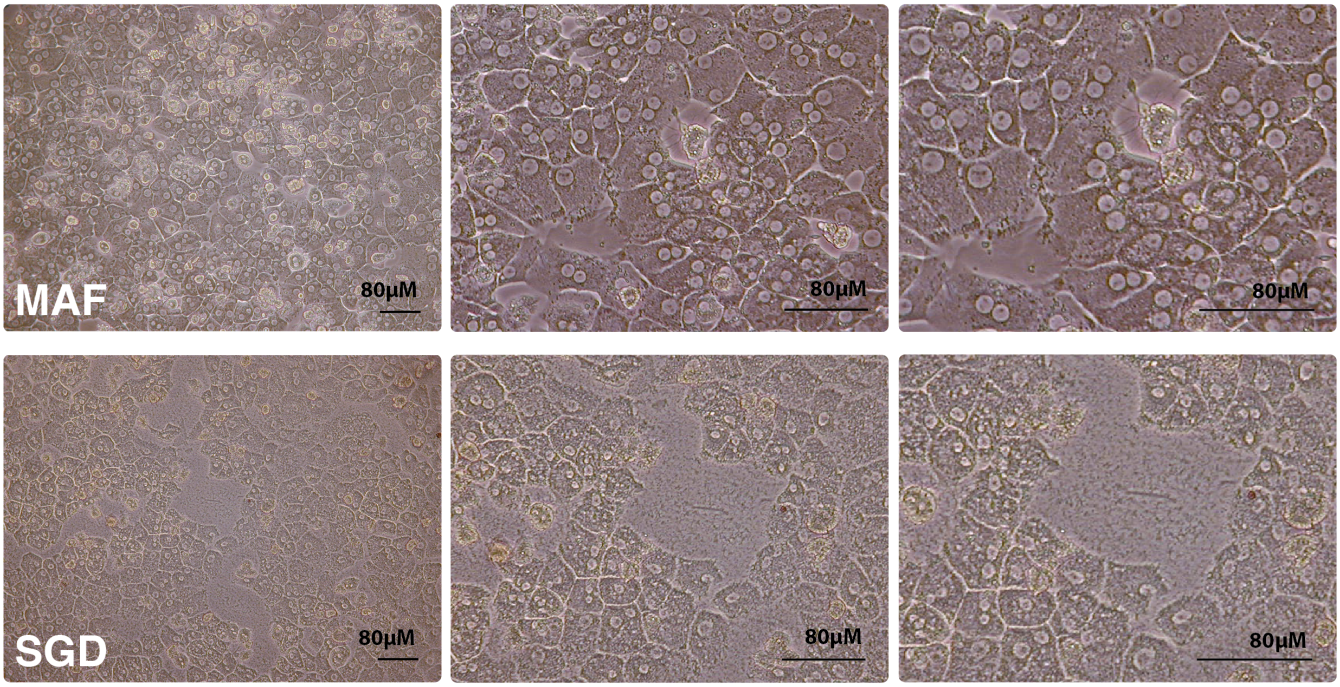


**Figure S6 | Sporozoite-associated morphological changes in primary rat hepatocytes.** Brightfield images of primary rat hepatocytes 20 hr after addition of sporozoites obtained by either MAF (top row) or SGD (bottom row). Cultured in 1% P/S.

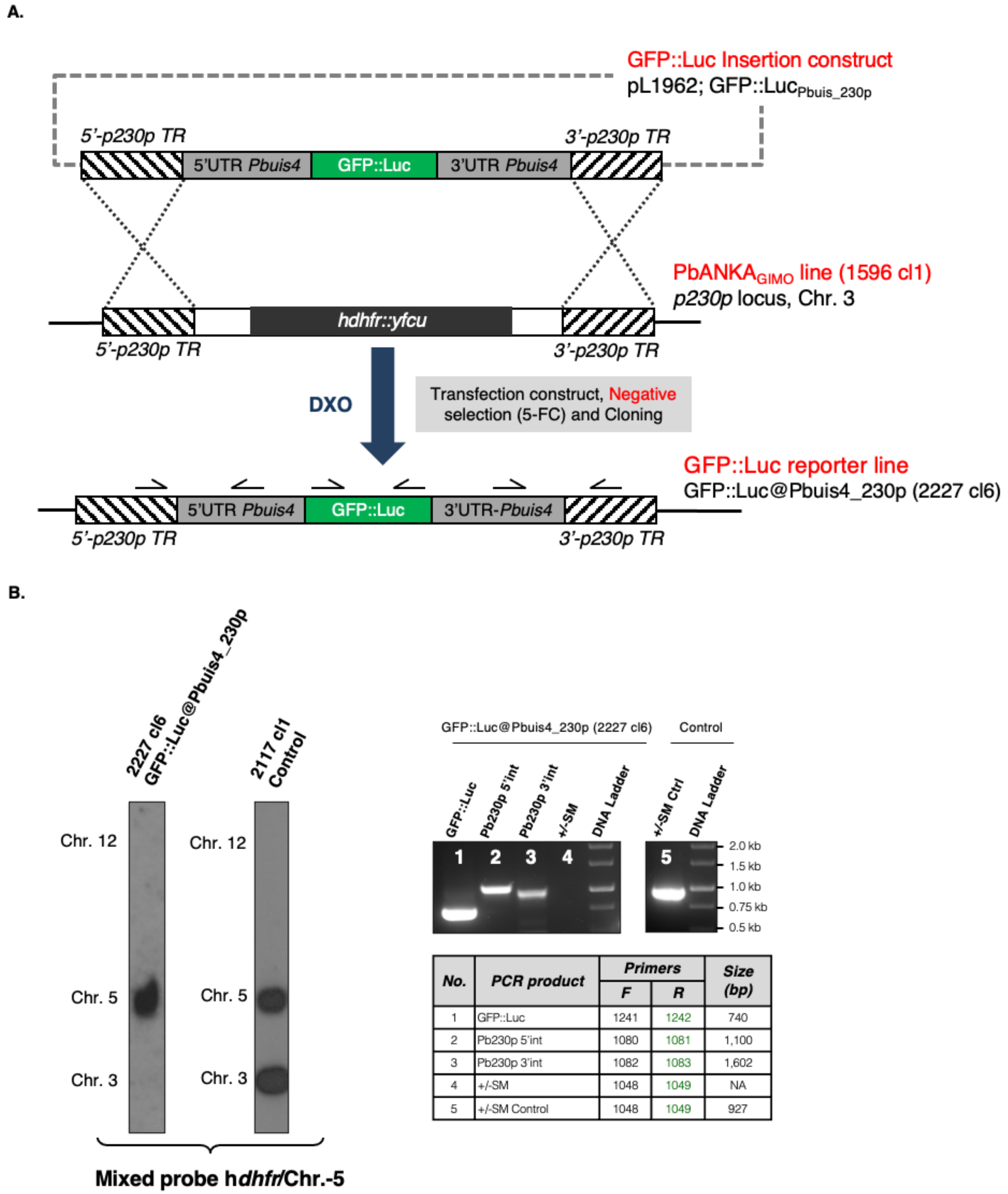


| No. | PCR product | Primers | | Size (bp) |
|---|---|---|---|---|
| | | F | R | |
| 1 | GFP::Luc | 1241 | 1242 | 740 |
| 2 | Pb230p 5'int | 1080 | 1081 | 1,100 |
| 3 | Pb230p 3'int | 1082 | 1083 | 1,602 |
| 4 | +/-SM | 1048 | 1049 | NA |
| 5 | +/-SM Control | 1048 | 1049 | 927 |

**Figure S7 | Generation and genotype analysis of the reporter line *PbANKA*-GFP::Luc@Pbuis4_230p.** A) Schematic representation of the introduction of the GFP::Luciferase expression cassette into the genome of the GIMO PbANKA parent line 1596 cl1. The *gfp-luciferase* fusion gene is under the control of *Pbuis4* regulatory sequences (5'UTR and 3'UTR regions). DNA construct pL1962 containing the GFP::Luciferase expression cassette is integrated into the modified *P.*

*berghei p230p* locus on chromosome (chr.) 3, containing the h*dhfr*::y*fcu* selectable marker (SM) cassette (black box), by double cross-over homologous recombination (DXO) at the *p230p* target regions (hatched boxes). Negative selection with 5-FC selects for parasites (line 2227 cl6) that have the GFP::Luciferase expression cassette introduced into the neutral *p230p* gene locus and the h*dhfr*::y*fcu* marker removed. Location of primers used for PCR analysis and sizes of PCR products are shown. See Supplementary Table 1 and 2 for primer sequences. B) Conformation of correct integration of the GFP::Luciferase expression cassette into the genome by Southern analysis of Pulsed Field Gel (PFG) separated chromosomes and diagnostic PCR. Left panel: Hybridisation of PFG separated chromosomes (chr.) of the reporter line *PbANKA*-GFP::Luc@Pbuis4_230p A) shows integration of the expression cassette into the *p230p* GIMO locus on chromosome (chr.) 3 by the absence of the h*dhfr::*y*fcu* SM cassette in the cloned reporter line. The Southern blot was hybridized with a mixture of two probes: one recognizing h*dhfr* and a control probe recognizing chr. 5. As an control parasite line, line 2117 cl1 was used with the h*dhfr::*y*fcu* SM integrated into chr. 3. Right panel: Diagnostic PCR shows the absence of the h*dhfr*::y*fcu* SM, the presence of GFP::Luciferase gene and the correct integration of the expression cassette into the genome of *PbANKA*-GFP::Luc@Pbuis4_230p at the 5'- and 3'-regions of *p230p* (5'int and 3'int). See **A** for the primers' locations and Supplementary Table 2 for primer sequences.

**Table S1 | Primers for generation of pL1962 DNA construct**

| DNA Construct | Primer No. | Primer Sequences * | Restriction Sites | Fragment Size (bp) | Description |
|---|---|---|---|---|---|
| **pL1962** | **7169** | tat**cctgcagg**GTGATAGTGTAGATTTTTTTGTTTGAC | SbfI | 1,519 | Pbuis4 5'-UTR promoter sequence, F |
| | **7170** | ataagaat**gcggccgc**AGACGTAATAATTATGTGCTGAAAGG | NotI | | Pbuis4 5'-UTR promoter sequence, R |
| | **7171** | cg**gatatc**TATAATTCATTATGAGTAGTGTAATTCAG | EcoRV | 1,025 | Pbuis4 3'-UTR sequence, F |
| | **7172** | ggcc**ggtacc**TTTCGCTTTAATGCTTGTCATC | KpnI | | Pbuis4 3'-UTR sequence, R |
| | **7295** | ataagaat**gcggccgc**GATCTATGAGTAAAGGAGAAGAAC | NotI | 2,448 | GFP::Luc, F |
| | **7296** | CTAGAATTACACGGCGATCTTTCC | -- | | GFP::Luc, R |

* Red color: Restriction site sequence

**Table S2 | Primers for genotyping the GFP::Luc@Pbuis4_230p (2227 cl6) reporter line**

| Primer No. | Description | Primer sequence |
|---|---|---|
| **1048** | hDHFR-yFCU (+/-SM) F | ATCATGCAAGACTTTGAAAGTGAC |
| **1049** | hDHFR-yFCU (+/-SM) R | CATCGATTCACCAGCTCTGAC |
| **1080** | 5'-*p230p* Integration F | ACTGTTATATTTGGTGATGGAATGG |
| **1081** | 5'-*p230p* Integration R | TATACATCCACGGATGCATAGAAG |
| **1082** | 3'-*p230p* Integration F | TCTGCATTAACCTTAAATATGAAAAACAC |
| **1083** | 3'-*p230p* Integration R | TTCAGTGAAATCGCAAACATAAGTATC |
| **1241** | GFP::Luc F | ccgg**ggtacc****ctcgag**ATGAGTAAAGGAGAAGAACTTTTCACTG |
| **1242** | GFP::Luc R | cgA**GGATCC**CTTTGTATAGTTCATCCATG |

* Red/blue color: Restriction site sequence
